# A dynamic Boolean model of molecular and cellular interactions represents psoriasis development and predicts drug candidates

**DOI:** 10.1101/2023.09.03.556147

**Authors:** Eirini Tsirvouli, Vincent Noël, Åsmund Flobak, Laurence Calzone, Martin Kuiper

## Abstract

Psoriasis is a chronic skin disease affecting 2-3% of the global population. Psoriasis arises from complex interactions between keratinocytes and immune cells, leading to uncontrolled inflammation, immune hyperactivation and perturbed keratinocyte life cycle. Although the latest generation of drugs have greatly improved psoriasis management, the disease remains incurable, and the substantial variability in treatment response calls for novel approaches to comprehend the intricate mechanisms underlying disease development and to discover potential drug targets. In this study, we present a multiscale population model that captures the dynamics of cell-specific phenotypes in psoriasis, integrating discrete logical formalism and population dynamics simulations. Through simulations and network metrics, we identify potential pairwise interventions as alternative treatment options. Specifically, The model predictions suggest that targeting neutrophil activation in conjunction with either PGE2 production or STAT3 signaling shows promise comparable to IL-17 inhibition, which is currently the most used treatment option for moderate and severe cases of psoriasis. Our findings underscore the significance of considering complex intercellular interactions and intracellular signaling cascades in psoriasis, and highlight the importance of computational approaches in unraveling complex biological systems for drug target identification.

**Author summary:** In our study, we aimed to uncover the complex mechanisms underlying psoriasis and identify potential treatment options. By utilizing a computational model, we simulated the dynamic interactions between different cell types involved in psoriasis, such as immune cells and keratinocytes. Our model predicts that targeting neutrophil activation, combined with either PGE2 production or STAT3 signaling, may yield comparable effectiveness to the current standard treatment for moderate or severe psoriasis, namely IL-17 inhibition. Our study underscores the importance of computational modeling in unraveling the complexities of disease systems and provides a foundation for identifying new candidate treatment options in psoriasis that should be tested in the lab.

## Introduction

Psoriasis is a chronic skin disease, characterized by a dysregulation of inflammation and immune processes, that is estimated to affect 2-3% of the global population. The full etiology of psoriasis is multifactorial, with genetic predisposition and environmental factors modulating disease susceptibility, therapy response, and disease course [1]. Psoriasis can be described as a complex, dysregulated multicellular system in which the interaction between immune cells and keratinocytes results in the emergence of the disease’s various phenotypes. The main hallmarks of psoriasis include immune activation and keratinocyte hyperproliferation, which involve the activation of inflammatory cascades and the release of cytokines, chemokines, and growth factors [2]. These factors create a perpetual loop that sustains and exacerbates immune responses while promoting epidermal hyperproliferation. Due to the complex nature of psoriasis, a multiscale systems approach is expected to aid understanding of its pathophysiology and therapeutic options.

Psoriasis evolves in two main stages: the initiation stage and the maintenance stage. Both stages are known to involve multiple positive feedback loops of signaling between keratinocytes and immune cells. Initially, keratinocytes can respond to a variety of inflammatory triggers that can be either external (e.g., skin injury or drugs) or internal (e.g., infections or mental stress) [1]. In response to such triggers, keratinocytes release pro-inflammatory mediators that promote immune cell activation and infiltration into the skin [3]. Such mediators include the antimicrobial peptide LL-37 that, in complex with either DNA or RNA, recruits and activates plasmacytoid dendritic cells (pDCs). Active pDCs produce high levels of interferon (IFN)-α that activate myeloid DCs (mDCs). Mature mDCs shape the differentiation landscape of naive T-helper (Th) cells towards the Th1, Th17, and Th22 subtypes [4–6]. These subtypes and their secreted cytokines dominate the cytokine microenvironment, and keratinocytes are considered the main responders to Th cell-derived cytokines [7]. More specifically, IFN-γ and tumor necrosis factor alpha (TNFα) are mainly produced by Th1 cells, whereas interleukin-17 (IL-17) is produced by Th17 cells, and IL-22 is produced by both Th17 and Th22 subtypes [7]. Keratinocytes stimulated by these cytokines have a disrupted life cycle characterized by unabated proliferation, aberrant differentiation, and reduced apoptosis. Additionally, psoriatic keratinocytes adopt an inflammatory phenotype and secrete to their environment additional cytokines and chemokines, which promote the survival and proliferation of immune cells and further recruit additional cell types, such as neutrophils, thereby sustaining chronic inflammation [4]. The interplay between keratinocytes and immune cells creates a system where reciprocal cause and effect sustains the psoriatic disease phenotype.

Despite the significant advancements made in identifying the causes of psoriasis and developing effective treatments that work for many patients, there are still challenges to overcome. The high variability in treatment responses among individual patients and the ongoing need for affordable medication due to the chronic nature of the disease demand the design of new therapies to effectively manage psoriasis. Current treatment options for psoriasis include topical creams, phototherapy, systemic medications, and biologic agents. Biologics are a relatively new class of drugs that target specific molecules in the immune system that play a key role in the development and progression of psoriasis, and these have revolutionized the treatment of psoriasis. Examples include TNFα inhibitors (e.g., etanercept, adalimumab), IL-17 inhibitors (e.g., secukinumab, ixekizumab), and IL-23 inhibitors (e.g., guselkumab, tildrakizumab). While these treatments can effectively reduce symptoms and improve the quality of life for many patients, they often come with significant side effects, and not all patients respond equally to the same treatment [8]. The high variance in treatment response highlights the need for personalized medicine approaches to tailor treatment options to individual patients based on their unique genetic, molecular, and clinical characteristics. Drug switching and combination treatment have been proposed as potential solutions for these patients [9,10].

Previously, we have modeled the effects of IL-17 signaling in PGE2 production in psoriatic keratinocytes and are working on an extended version of this model to capture heterogeneous manifestations of the disease and differential responses to treatments when coupled with patient gene expression data (Preprint at [11]). Here, we present a multiscale population model which allows us to follow the dynamics of cell-specific phenotypes in a time-resolved manner. The model represents the key intercellular interactions and intracellular signaling cascades involved in the disease and is based on a discrete logical formalism and population dynamics simulated with UPMaBoSS [12,13]. Using network analysis through propagation metrics, we identified pairwise interventions that could serve as alternative treatment options and might be experimentally tested. Among those combinations, the targeting of neutrophils together with PGE2 production or STAT3 was predicted to have an effect similar to IL-17 inhibition.

## Results

### The PsoriaSys model - A multiscale model of psoriasis

The PsoriaSys model consists of 87 nodes and 235 edges. The model’s nodes can represent a cell type (e.g., neutrophils), a cell in a specific state (e.g., proliferating keratinocytes), a ligand (e.g., IL-17) or its receptors (e.g., IL-17R), a transcription factor (e.g., STAT3), or other intermediate signaling proteins. The model encompasses 18 cell nodes, and it describes the main intercellular interactions and intracellular signaling during the various psoriasis stages. The edges in the model represent the causal (activating or inhibiting) interactions between signaling components or transitions between different cell states.

UPMaBoSS is an extension of MaBoSS [14] that allows simulations of cell population dynamics. In this framework, cells can die, divide and interact. The size of a cell population is defined by multiplying the probability associated with an active cell node and the overall population size, for the same time point. The overall population size is calculated based on the *Division* and *Death* nodes, reflecting the increase or decrease of cell populations relevant to *t*=0. In the PsoriaSys model, the keratinocyte is the only cell type that influences the Division and Death nodes, and therefore only the proliferation and death of keratinocytes are captured by the model. Considering that keratinocytes comprise the cell population whose phenotype is most affected by disease progression and alleviation, the population levels of keratinocytes and their physiological state (proliferation versus differentiation) were used as a proxy for the disease course. In addition to proliferation, four more physiological states of the normal and diseased keratinocyte life cycle are represented: the normal, unstimulated keratinocyte (KC node), which ends its life cycle as a fully differentiated keratinocyte (Diff_KC node); the activated keratinocyte that responds to external or internal stimuli and initiates an inflammatory cascade (aKC node); the proliferating keratinocyte (Prol_KC) characteristic of the disease; and lastly, the pre-differentiated keratinocyte (preDiff_KC node), a state of psoriatic keratinocyte in which the cell is unable to differentiate fully. Other stages of the keratinocyte life cycle, such as desquamation, are not included in the model, as keratinocytes in these states are not affected by active immune cells and need not to be taken into consideration for this multiscale model [15].

To ensure that the model can recapitulate known behavior, the sequence of the main events during disease development was confirmed (Figure 1C). As it can be observed from the activation trajectories of the nodes, the model successfully captures the initial responses of KCs to psoriatic triggers that lead to the activation of dendritic cells. Activated dendritic cells then translocate to the lymph node to activate Th naive cells. The time needed for this translocation is encoded as the parameter associated with the activation rate of Th0 cells and captured as a small delay between DC activation and Th activation and differentiation. When in the lymph, secretion of cytokines from DCs shapes the differentiation of Th cells to Th1, Th17, and Th22 subtypes while reducing their ratio to their anti-inflammatory counterpart, Treg cells. Differentiated Th cells then infiltrate the skin and disrupt the KC life cycle by secreting cytokines that promote their proliferation, disrupt their terminal differentiation, and prevent their death. Proliferating KCs secrete cytokines and chemokines that further recruit and promote the survival of the various immune cells. In the meantime, cell types that link innate and adaptive immunity and further enhance inflammatory responses, such as M1 macrophages and neutrophils, are also activated.

**Figure 1.**
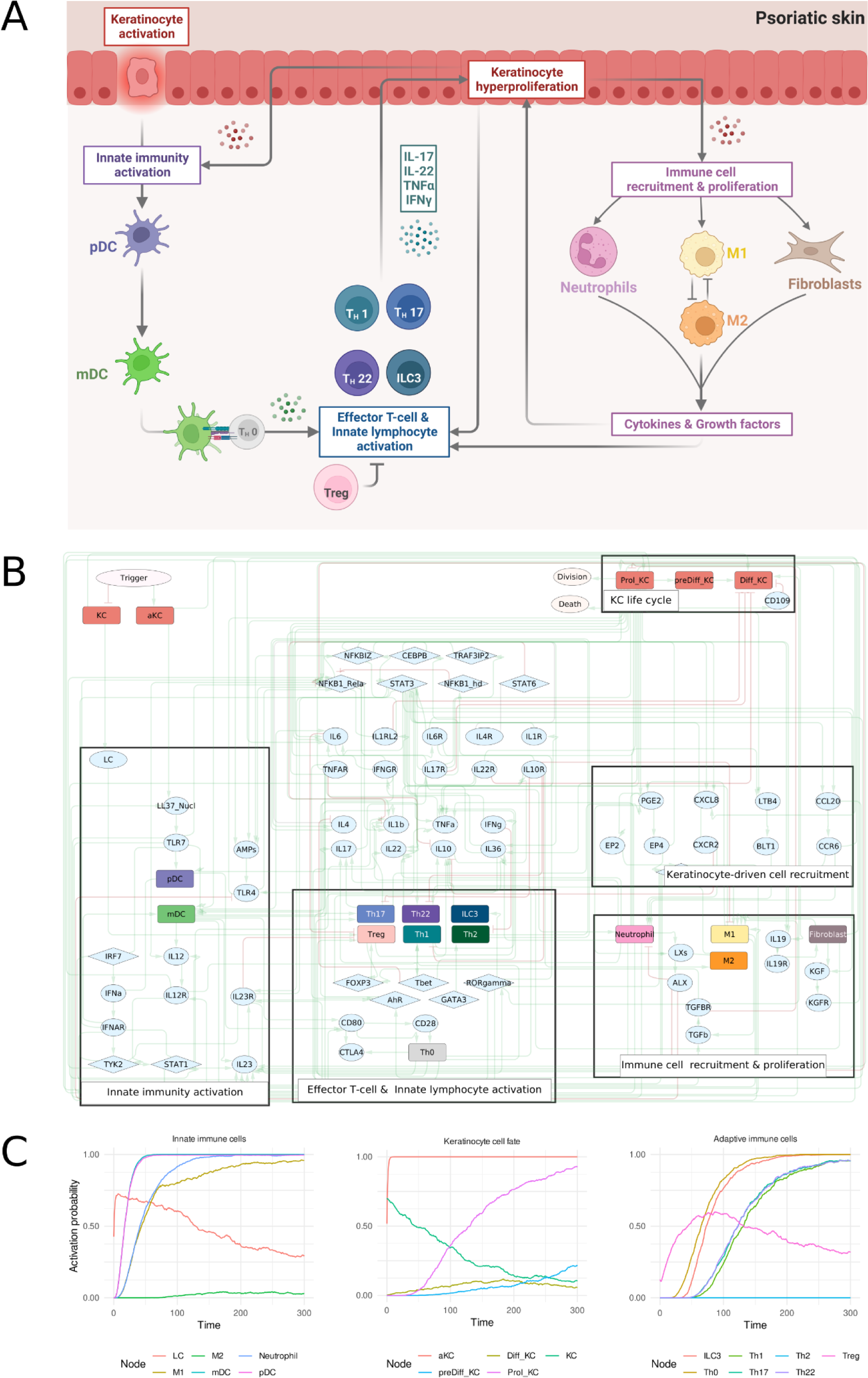
A. Schematic representation of the main cell types and interactions represented in the psoriasis model. pDC = plasmacytoid dendritic cells, mDC = myeloid dendritic cells, IL = Interleukin, Th = Thelper cells, Treg = T regulatory cells. B. The PsoriaSys model. Nodes represent cell types, or molecular entities. The nodes are colored based on the type of entity they represent (*see* figure legend). Green edges represent activating interactions and red edges represent inhibiting interactions. The diamond node represents the trigger of the disease. C. Sequence of events in response to the psoriatic trigger. At Time=0 a psoriatic trigger is applied. Abbreviations: LC = Langerhan cell, M1 = M1 macrophage, M2 = M2 macrophage, pDC = plasmacytoid dendritic cell, mDC = myeloid dendritic cell, KC = normal keratinocyte, aKC = trigger-activated keratinocyte, Prol_KC = proliferating keratinocyte, preDiff_KC = aberrantly differentiated keratinocyte, Diff_KC = terminally differentiated keratinocyte, ILC3 = Type 3 innate lymphoid cell, Th0 = naive T helper cell, Th = T helper cell, Treg = regulatory T cell.

### Sensitivity analysis

UPMaBoSS separates the rates of activation and inactivation of each node of the model, providing a means to modulate the timing of events, the behavior, and the long-term activity states of the system. A sensitivity analysis was conducted to evaluate the extent to which these parameters influence the system’s behavior and whether they have any effect on its outcome. The PsoriaSys model contains 155 parameters that determine each node’s activation or inactivation rate. These rates define how quickly an entity will be activated or inactivated during simulations. When available, the parameters were based on observations in the literature. As this type of information is scarce, only 17 parameters could be defined based on prior knowledge (Supplementary Table 1). All other parameters were set to match generally accepted principles and chosen to be the same for nodes of the same type. For example, cytokine and chemokine ligands were set to undergo rapid degradation/inactivation (i.e., high inactivation rate), resulting from extracellular signal depletion and other mechanisms of inflammation control [16]. For receptors, the activation rates were updated in each step of the simulations based on the probability of their ligand being active; the more the ligand is produced, the higher the activation rate of its receptor. To test the robustness of the model, a sensitivity analysis was performed by creating versions of the model where each of the parameters is either reduced or increased by 50% of their initial value, and the effects of these parameter changes were observed for the first 200 simulated hours following a psoriatic trigger. The deviation of the activation probability in the different model versions from the activation probability in the “wild type” (WT) model was used as a metric of the effect of a parameter on the model’s behavior. In a robust model, these deviations should be small and have a limited effect on the node’s state trajectory.

For most of the nodes, the changes in the model’s parameters do not significantly affect the behavior of the model, with almost all changes in the nodes’ activation probabilities deviating less than 0.1 from the WT probability (Supplementary Figure 1). While these deviations remained small, the state trajectories of a few nodes are sensitive to changes of certain parameters. The Treg (i.e. Regulatory T cells) node was sensitive to changes in the activation rates of some of its direct regulators; namely the increased activation rate of RORγ and AhR transcription factors and the reduced inactivation rates of CTLA4 and IL-12. RORγ and AhR are transcription factors related to T-helper cell differentiation towards Th17 and Th22. AhR activity suppresses Treg differentiation [17], and Th17 differentiation, mostly driven by RORγ, is mutually exclusive with Treg differentiation [18]. Furthermore, the activation probability of the dendritic cell nodes (i.e., mDC and pDC nodes) was sensitive to the activation rate of the LL-37/Nucleic acid complex, which is responsible for their activation in response to a psoriatic trigger. Lastly, the probability of active fibroblasts was affected by the reduced activation rate of proliferating keratinocytes affecting the probability of active fibroblasts. The relationship between fibroblasts and proliferating keratinocytes can be traced to the activation of fibroblasts by the KC-derived IL-19, which in turn stimulates keratinocyte proliferation by priming fibroblast to secrete keratinocyte growth factor (KGF) [19]. Comparing the size of the proliferating keratinocyte population between conditions shows that the activation and division of proliferating keratinocytes was not sensitive to changes in any of the model’s parameters, with this population increasing in size at a similar rate as in the “wild type” model. This behavior may result from how the model was built to represent the events of disease progression in the presence of psoriatic stimuli, with an active disease phenotype expected to be reached unless a mitigating perturbation is applied.

### Model validation

Having established that the model can show cell population dynamics consistent with the literature on disease progression (see above), a set of *in silico* perturbations was checked against known observations from a collection of physiological observations under certain conditions, such as subtypes of the disease or in response to treatment.

The effects of the most frequent treatment options were also validated by testing the response of the system to the inhibition of IL-17, TNFα, IL-12, and IL-23, all of which are the main targets of the biologics used to treat moderate-to-severe psoriasis [8]. The effect of treatment was simulated by using ‘chained’ simulations (Figure 2A). First, a trigger is applied to the system, encoded as the activation of the “*Trigger”* node. Upon this trigger, the system reaches an active psoriatic state. After this psoriatic state is reached, a second simulation is performed, in which an *in silico* perturbation is applied on the psoriatic system state that resulted from the initial trigger. In case of inhibitory perturbations the state of the perturbation target was fixed at 0, while in activating perturbations the target’s state was fixed at 1. The treatment effect is then compared to the predicted trajectory of the individual cell types where no treatment is applied.

**Figure 2.**
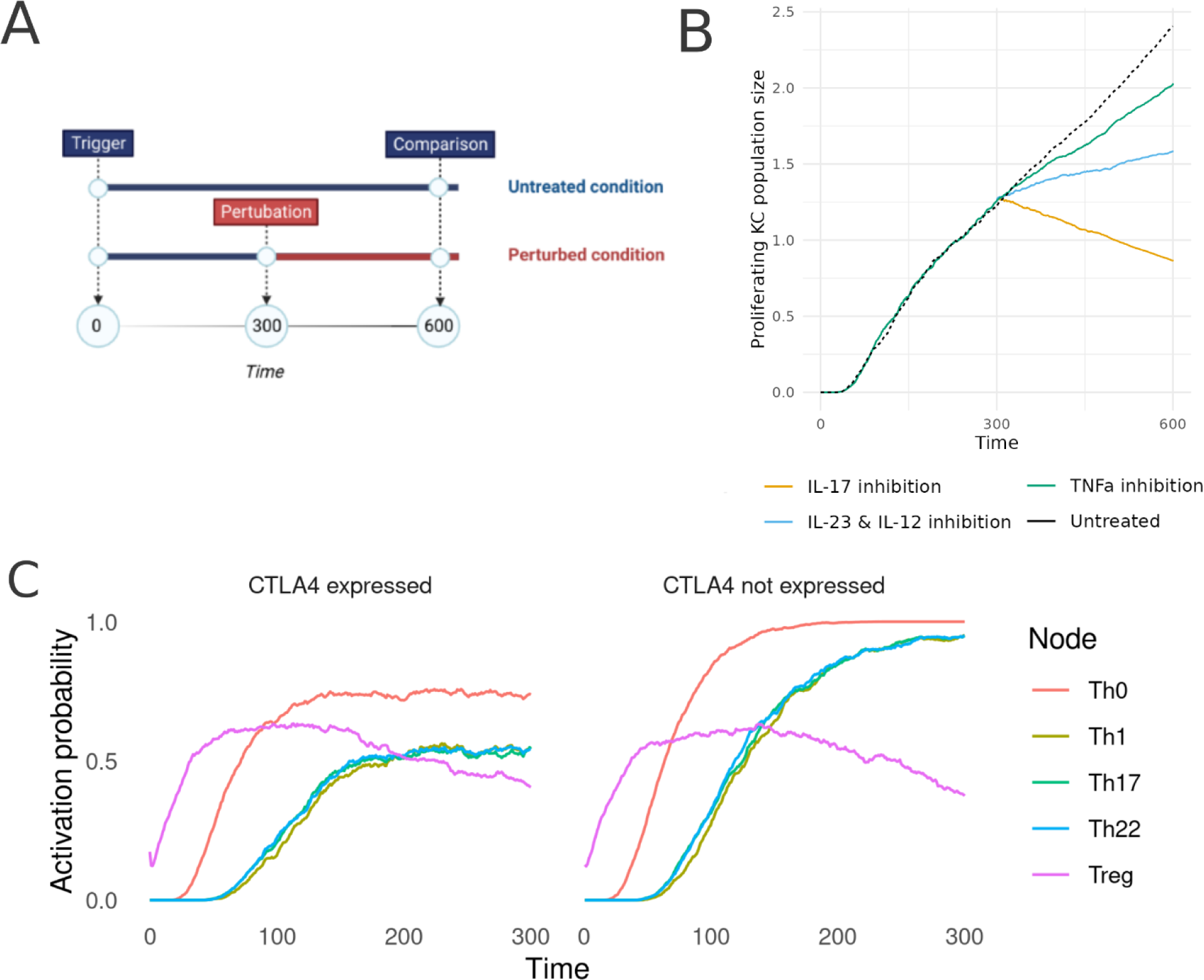
A. The effect of biologics in psoriasis on the population size of proliferating keratinocytes compared to the population size in untreated conditions. The x-axis represents time and the y-axis represents the relative size population of proliferating keratinocytes. B. Comparison of T-helper activation when T-helper cells express CTLA4 (left panel) and T-helper activation when CTLA4 is not expressed (right panel). The x-axis represents time, and the y-axis represents the probability of activation of each node.

Current biologic treatments mostly target the IL-23/IL-17 pathogenic axis in different stages of the disease development. IL-17 is a multifunctional cytokine widely characterized as a key driver of psoriasis by regulating cell fate proliferation and differentiation and sustaining the release of inflammation-promoting agents, such as antimicrobial peptides [3,20]. Biologics targeting IL-17, for instance ixekizumab and brodalumab, have shown an excellent efficacy in treating moderate-to-severe psoriasis and are relatively well-tolerated with long-term maintenance of treatment responses [21]. Targeting IL-17 in the multiPSO model leads to the biggest reduction of proliferating KC population (∼ 64% decrease) compared to the simulated single-target treatments (Figure 2B & Supplementary Table 3). However, IL-17 inhibition has a relatively limited effect on the rest of the cell types (Supplementary Table 3). As keratinocytes and generally non-hematopoietic cells are the main targets of IL-17 [20,22], it was hypothesized that by inhibiting IL-17, the normal KC life cycle is restored, which also leads to the limited production of inflammation-inducing agents and the subsequent reduction of other cell types, that could be considered as a secondary effect.

TNFα is involved in the earlier events of psoriasis and drives the induction of dendritic cells and their maturation [23]. Additionally, TNFα acts synergistically with IL-17 on KC responses, enhancing the expression of main inflammatory drivers, KC-derived cytokines, and other immunomodulating molecules [24]. Several TNF inhibitors, such as adalimumab and etanercept, have been approved for treating moderate-to-severe psoriasis; however, all with a variable efficiency among different patients [25]. The inhibition of TNFα can induce various responses in patients, including the induction of paradoxical psoriasis, a type of psoriasis that is Th cell-independent and driven by Type-I IFN signaling [23,26]. The inhibition of TNFα in the PsoriaSys model results in a 15% reduction in the KC population (Figure 2B & Supplementary Table 3). In addition to its effect on KC, TNFα amplifies inflammatory responses in several ways. Fibroblasts stimulated with TNFα and IL-8 are not able to produce IL-10, a potent anti-inflammatory cytokine [27]. Additionally, IL-10 production in response to TNF blockade has been observed *in vitro* [28]. With the reduction of proliferating KC and subsequently the KC-derived IL-8, fibroblasts secrete IL-10 activating anti-inflammatory M2 macrophages. The production of IL10 leads to the eventual repression of Th22 cells and, therefore, lower IL-22 levels. According to the model, TNF blockade reduces the KC population by inducing their death. *In vitro* experiments on human epidermal keratinocytes (HEKs) showed that IL-22 plays a strong antiapoptotic role in the cell fate of KC, independently of IL-17 [29]. This mechanism could explain the reduced population of proliferating keratinocytes.

Lastly, the effect of inhibiting IL-12 and IL-23 was tested. Both cytokines promote T-cell-mediated responses by shaping the differentiation of Th cells towards the Th1 and Th17 phenotypes. Ustekinumab is an antibody that blocks IL-12 and IL-23 signaling by binding and neutralizing the shared p40 subunits of the two cytokines [30]. IL-12/IL-23 inhibition reduces the KC population by 34% (Figure 2B & Supplementary Table 3) and completely inhibits Th cell activation and differentiation. Comparative studies between biologics targeting IL-17, IL-23, TNF, and IL-12/IL-23 showed that IL-17 and IL-23 inhibitors have a higher efficiency than IL-12/IL-23 and TNF inhibitors [31]. The model successfully predicts a higher effect of IL-17 inhibition compared to TNF inhibitors, but IL-23 and the combinatorial inhibition of IL-12 and IL-23 shows similar levels of effect on KC populations. While the model can be used to compare the probabilities of a cell being activated or inhibited between different conditions, other more quantitative models, such as pharmacodynamics/pharmacokinetics models, could be used to quantify treatment effects in more detail.

In addition to treatment responses, we also found that the model could distinguish between disease development of mild and more severe cases of psoriasis. In the study of [32], CTLA4 was proposed to be upregulated in patients with milder psoriasis. T-cell activation is regulated by co-stimulatory molecules that can either promote or inhibit their activation. CD28 is a receptor in T naive cells that binds to CD80 and CD86 ligands on the surface of antigen-presenting cells (such as DCs). CTLA4 is another co-stimulatory molecule on the surface of T cells similar to CD28, which competitively binds the CD80/CD86 ligands with a much higher affinity than CD28. Contrary to CD80, CTLA4 elicits an inhibitory signal for T-cell activation. After the activation of T cells by CD80/86-CD28 interactions, CD28 is internalized and degraded, and CTLA4 translocates to the cell surface. This negative regulation provides a control mechanism to regulate T-cell activation and inflammation, which does not occur in psoriasis. The effect of CTLA4 was encoded in the regulation of the Th0 node by increasing the rate of its downregulation when CTLA4 is expressed. The development of the disease in two model instances, one with a functioning CTLA4 and one without, shows that CTLA4 expression can prevent full activation of Th cells, potentially leading to a milder manifestation of the disease, as shown in Figure 2C.

### Perturbation analysis

Having confirmed that the model adequately represents the basic regulatory network underlying psoriatic development, a systematic perturbation analysis was performed to identify new potential drug intervention points. While targeting each druggable component, the inhibition of inflammatory components and the activation of anti-inflammatory components were tested. The effect of each perturbation was assessed by model readouts that mirror measurements commonly used in a clinical setting to assess treatment response, namely reduction of skin thickening (represented as the proliferating KC population size in the model), T-helper and Treg activation, and immune cell recruitment. For each of the treatments, the response was quantified as the change in probability of a node to be activated by the formula:

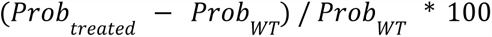

The responses to the different treatments were highly variable. The results of all tested perturbation can be found in Supplementary Table 3. In addition to the effectiveness of the known biologics (Figure 2B), the inhibition of TYK2 was among the most effective perturbations concerning the afore-described criteria. TYK2 inhibition is predicted to not only reduce KC proliferation (by 33%) and abolish the activation of Th cells. TYK2 is part of the JAK-STAT signaling pathway that mediates the expression of several drivers of psoriasis, such as Type I IFNs, IL-12, and IL-23, making TYK2 an integral part of inflammation and immune response regulation. A similar effect with the TYK2 inhibition is predicted for other components of the IL-23/IL-17 axis, such as the inhibition of IL-23 and STAT3. However, the differentiation of the Th1 cells is not inhibited, as IL-12 expression, the driver of Th1 differentiation, is not affected in any way by targeting the IL-23/IL-17 axis.

Another promising target proposed by the model is prostaglandin E2 (PGE2), whose inhibition reduces the size of the proliferating KC population and inhibits Th cell activation. PGE2 is a lipid mediator produced by arachidonic acid (AA) and elicits a wide range of biological effects in inflammation and its resolution. PGE2 acts through four main Prostaglandin E2 receptors (EP1-EP4), all of which are connected to a diverse set of downstream pathways. EP2 and EP4 are upregulated in psoriasis [33,34] and are the primary receptors through which PGE2 exerts its pathogenic effect in psoriasis. Due to their partial cross-talk, the inhibition of either EP2 or EP4 does not significantly reduce KC proliferation. However, EP4 inhibition shows a mild effect on the reduction of early activated immune cells (e.g., inflammatory DCs), naive Th cell activation, and, ultimately, the complete inhibition of Th17. An activating perturbation of IL-10 was among the few tested perturbations which, in addition to Th cell inhibition, leads to the activation of Treg cells. Additionally, it results in a phenotype shift of macrophages from the pro-inflammatory M1 phenotype to the anti-inflammatory M2 phenotype, and a reduction of neutrophil activation. However, it shows no effect on the KC populations. This observation corroborates the results of clinical trials on IL-10 therapeutics, where moderate improvement of psoriasis severity was attributed to the modulation of immune cell activation, with no observed direct effect on keratinocytes [35].

### Identification of high-influence node sets for combinatorial perturbation

After identifying effective single-node perturbations, the model was used to predict the effectiveness of combinatorial perturbations. As psoriasis is characterized by multiple feedback loops that amplify and sustain the disease’s phenotypes [4,36], it was hypothesized that the targeting of entities that are part of those feedback loops might restore normal phenotypes. A Feedback Vertex Set (FVS) is a set of nodes that includes at least one node from each feedback loop of a model, and in the PsoriaSys model 27 nodes (∼34% of the total nodes) are part of at least one feedback loop. However, due to the size of the FVS, targeting all of its components would not be a feasible option in experimental or clinical settings. To overcome this challenge, network propagation metrics were used to identify subsets of the complete FVS (also called minimum FVS) that could potentially serve as drug targets against psoriasis. Minimum FVS of two nodes were selected, based on the fact that pairwise combinations are the most commonly investigated in clinical settings. These subsets were ranked based on the overlap of the propagation metrics as described in [37]. A higher ranking of a node pair indicates a greater expected influence on the model’s states when the pair is perturbed.

In total, 333 two-node FVS were identified, and 62 of those pairs included non-druggable nodes. Druggable nodes represent transcription factors, ligands, or receptors, contrary to nodes representing cell types (e.g., Th0) or cell states (e.g., proliferating keratinocytes). Among the 217 druggable target pairs, cytokines, which are considered drivers of the disease and are already targets of the standard therapy options, were dominating. The top 20 target pairs (Supplementary Table 2) were further analyzed. If both a cytokine and its receptor appeared together with the same second target, only the cytokine inhibition was simulated as single-node perturbation analysis showed that inhibiting either a cytokine or its receptor had almost identical effects on the system.

Interestingly, the only cell nodes in the FVS represented proliferating keratinocytes, Th1, Th17, and neutrophils. The role of KC and Th cells in the self-sustained inflammatory cycle of psoriasis is well-described, therefore, suggesting that their reduction would reverse psoriatic phenotypes. While neutrophil recruitment is a histological characteristic of psoriasis, their pathogenic role in the disease remains unclear. To explore the potentials of neutrophils as a target we tested the effect of their inhibition with the PsoriaSys model. Neutrophils are one of the primary innate immune cells and are activated by a diverse set of inflammatory stimuli. Neutrophils can link innate and adaptive immunity by interacting with DCs and T cells and are attracted to psoriatic lesions by several chemokines released by KCs and T cells. On site, they induce respiratory burst, degranulation, and neutrophil extracellular traps (NETs) formation. Such processes can stimulate pDCs by activating TLR receptors which start the inflammatory cascade that leads to the pathogenesis of psoriasis. Additionally, neutrophils can amplify immune responses through several mechanisms, such as the release of proteinase-3 that cleaves pro-IL-36 to its activated form and the release of IL17, for which it is one of the main sources [38]. Among the top 20 target pairs proposed (Supplementary Table 2), neutrophils were implicated in 8 of them (Figure 3A). Notably, the combinatorial targeting of components of the IL-23/IL-17 axis along with neutrophil activation was ranked as the most influential target pairs. In the model, the role of neutrophils is restricted to their effect in psoriatic signaling and cytokine release, while processes such as the formation of neutrophil extracellular traps (NETs) are difficult to represent in the current modeling framework. The inhibition of the neutrophils alone has only a limited effect on the psoriatic phenotype, reducing proliferating keratinocytes by only 5% and no reduction in the main immune cells. However, a potential synergistic response is observed when neutrophil inhibition is combined with the inhibition of PGE2 or STAT3. The single inhibition of PGE2 or STAT3 reduces proliferating keratinocytes by ∼33%. When combining PGE2 and neutrophil inhibition, the population of proliferating keratinocytes is reduced to levels (63%) similar to the most effective single inhibition (i.e., IL-17 inhibition). Additionally, this combination restores the terminal differentiation of keratinocytes, as captured by their increased activation probability, and it reduces the probability of activation of the pre-differentiated populations. Furthermore, it completely abolishes the activation of all Th cell populations, ranking this combination among the most effective for restoring a normal phenotype from a psoriatic state. When the inhibition of neutrophils is combined with the inhibition of STAT3, we also notice a significant reduction of proliferating KCs (57%) (Figure 3B). For the other cell types, the changes are mostly STAT3-driven, and only a limited effect on Th1 activation or innate immune cells is observed.

**Figure 3.**
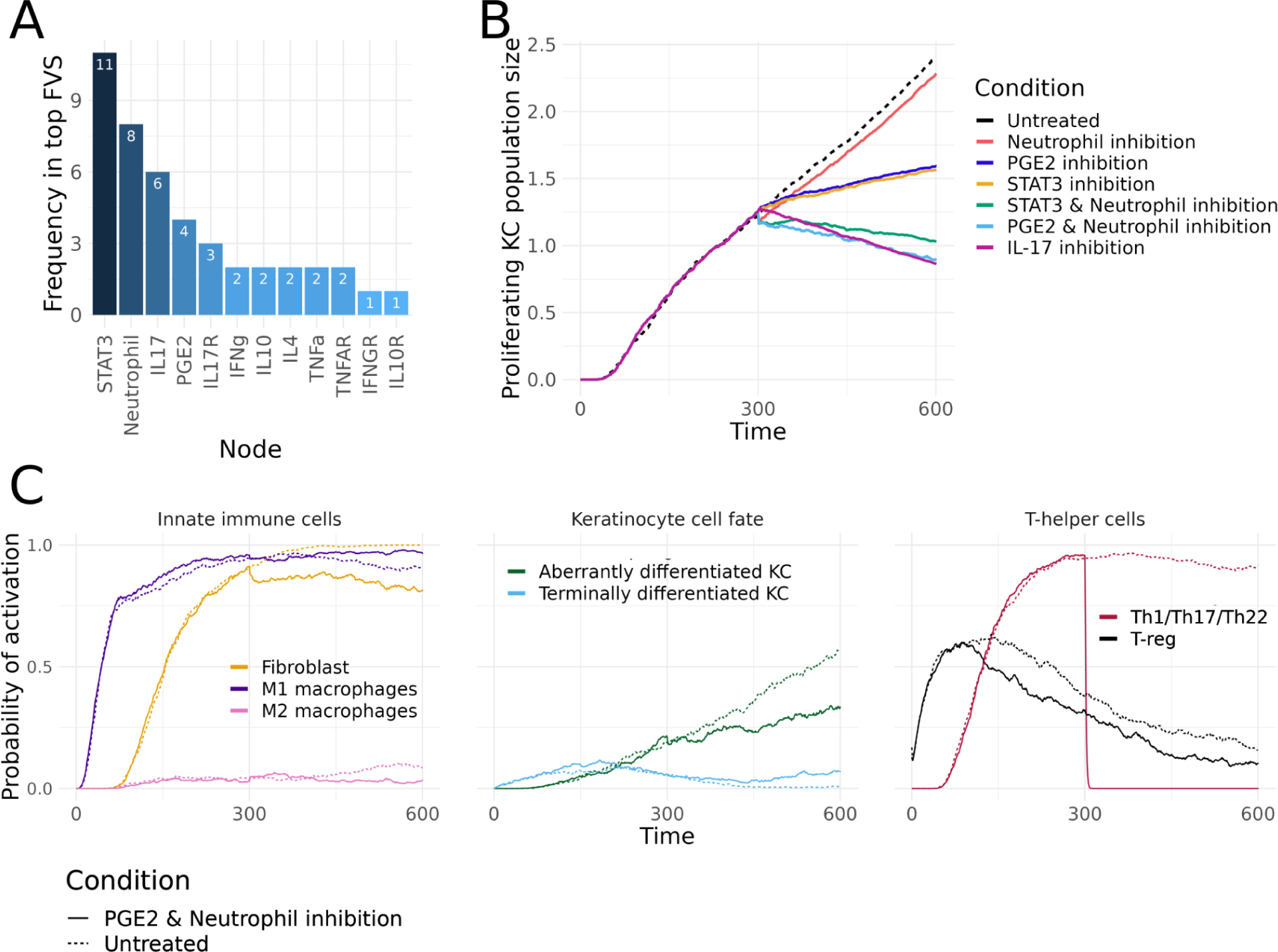
A. Frequency in which a node was included in minimum Feedback Vertex Sets (FVS), as identified by the intersection of three network propagation metrics (i.e., PRINCE, modifiedPRINCE and CheiRank). B. Comparison of the population size of proliferating keratinocytes (x-axis) for the combinatorial targeting of neutrophils and PGE2 production, or neutrophils and STAT3. The population sizes for untreated conditions (dashed line) and the inhibition of neutrophils, PGE2, and STAT3 alone are included for reference. C. Trajectory of the probability of activation of the cell nodes that respond to the combinatorial inhibition of neutrophils and PGE2 compared to their trajectories in untreated conditions (dashed lines). The y-axis shows the probability of activations and the x-axis shows Time.

Next on the ranked list of target pairs, the inhibition of IL17R and STAT3 was simulated. The combination of IL-17 and STAT3 leads to a similar reduction of proliferating keratinocytes as the inhibition of IL17 alone. However, it also reduces other immune cells, including M1 macrophages, fibroblasts, and all Th cells, and increases the activation of the anti-inflammatory Treg cells by 23%. STAT3 has a multifaceted role in psoriasis and influences several cell types. Activated by IL-23 in Th cells, STAT3 induces the expression of RORγ, which promotes Th17 responses and the expression of IL-17A, IL-17F, IL-22, and IL23R, further amplifying inflammatory responses. In keratinocytes, STAT3 is activated downstream of IL-17 and IL-22 responses [19,39] and produces CCL20 in an IL-1b-induced way [40]. Therefore, this double inhibition could inhibit psoriatic phenotypes by inhibiting the alternative ways the uncontrolled inflammation might arise. The results of other combinations with IL-17, namely neutrophil inhibition or IL-4 induction, did not differ from IL-17 single inhibition results.

For the remaining target pairs, the combinatorial perturbation of the IL-23/IL-17 axis and entities with a role in inflammation amplification were also highly ranked. These combinations include the double inhibition of STAT3 with either PGE2, TNFα, or IL-4 (Figures 3B and 3C). However, these combinations show limited, if any, additional effects compared with the single perturbations.

Similarly, none of the combinations that included IL-4, result in an effective reduction of psoriatic phenotypes. IL-4 is the only cytokine that shifts the differentiation of Th cells towards a Th2 phenotype [41]. The imbalance between Th1 and Th2 cytokines is associated with the development of psoriasis [42] and correlated with disease severity [43]. Shifting the T-cell differentiation towards Th2 by administering IL-4 has been shown to alleviate symptoms in patients with severe psoriasis [41]. However, the model could not replicate this observation, potentially suggesting some gaps in the knowledge of IL-4 driven mechanisms and therefore their underrepresentation in the model.

## Discussion

We have developed a logical, multiscale model of psoriasis (PsoriaSys model) that integrates molecular interactions, signal transduction, and cell-to-cell communication between psoriatic cells and their surrounding environment. The PsoriaSys model represents the biological events that occur upon the presence of psoriatic triggers that lead to the initiation of the disease and mechanisms and pathways driving disease development and maintenance. The model links the dysregulated signal transduction processes to the secretion of growth factors, cytokines and chemokines, and it includes intercellular communication between psoriatic skin cells, namely keratinocytes and immune cells.

This PsoriaSys model uses logical formalism combined with stochastic simulations. Logical formalism allows the discrete modeling of psoriatic interactions. In contrast, the stochastic simulations allow the exploration of temporal evolution beyond the strict state changes supported by this qualitative framework and the analysis of the probabilities of nodes to reach a specific state. This combination was valuable for replicating the development of psoriatic phenotypes in response to psoriatic triggers and emulating the known effect of drug perturbations. Additionally, model instances allow to capture the effects of some of the known sources of patient heterogeneity. This was showcased by simulating the effect of lack of CTLA4 expression, where the model recapitulates the effect of CTLA4 in immune cell activation. CTLA4 is a co-inhibitory molecule which acts by limiting T helper cell hyperactivation, and CTLA4 expression has been found to be inversely correlated with psoriasis severity [32,44]. Having validated the descriptive properties of the model, it was next used to predict the effect of perturbations of single entities or their combinations. It was shown that the model correctly identifies IL-17 inhibition as the most effective in reducing the population size of proliferating keratinocytes. In clinical settings, IL-17 blockade through biologics has provided a treatment option against moderate-to-severe and severe cases [21] with unprecedented efficacy. Among other effective perturbations, the inhibition of PGE2 or TYK2, and the activation of IL-10 were proposed. Recently, the first TYK2 inhibitor (i.e., deucravacitinib) was approved by the U.S. Food and Drug Administration (FDA) for the treatment of moderate-to-severe plaque psoriasis. The model’s ability to predict known targets highlights that the model accurately represents the biological regulatory wiring that drives the pathogenesis and development of psoriasis. The role of PGE2 is multifaceted in psoriasis, with the prostaglandin being involved in multiple aspects of disease development. Several studies show that PGE2 shifts adaptive immunity toward Th1 and Th17 responses by affecting DCs [45], priming the expansion of Th17 cells, and promoting the differentiation of Th1 cells [46]. The effect of PGE2 on cell fate decisions of KC has been previously described by our work on the computational and experimental modeling of KC behavior upon cytokine and PGE2 stimulation [33]. Inhibitors against cPLA2a, the phospholipase regulating PGE2 expression, have shown high efficacy against plaque psoriasis in clinical trials [47], and while several well-established psoriasis drugs are reducing EP and, subsequently, PGE2 levels, the direct inhibition of EP receptors has not been fully explored in psoriasis, but it is being investigated in other diseases, such as treatment of solid tumors [48].

Furthermore, using network propagation metrics, we identified pairs of model components whose combinatorial targeting was more likely to control the system’s behavior. The double inhibition of neutrophil action and PGE2 or STAT3 reduced psoriatic phenotypes to similar levels as the IL-17 blockade. Neutrophils act as immune regulators in psoriasis, bridging innate and adaptive responses [38], and their accumulation on psoriatic skin is considered a hallmark of psoriasis [49]. During degranulation and respiratory burst, neutrophils release, among others, granule-derived serine proteinases, such as NE, proteinase 3, and cathepsin G, which can activate by cleavage various psoriatic cytokines, such as TNFα and IL-36 [38]. Furthermore, neutrophil clearance is associated with successful treatments, for instance anti-IL-17 agents [38,49,50]. In general, anti-IL-17 drugs are hypothesized to act on neutrophil-derived IL-17 and disrupt neutrophil and keratinocyte crosstalk [22]. Based on the model, we suggest that neutrophils could serve as a promising target against psoriasis, either alone or in combination with other components of the IL-17/PGE2 axis components.

While the model recapitulates the known events of the disease, it will be interesting to challenge predictions of the model with new experimental data, however, this is beyond the scope of this work. As in every modeling effort, the model’s ability to represent specific events is limited by the available knowledge about the mechanisms underlying such events. Additionally, technologies such as single-cell RNA sequencing could aid with defining and fine-tuning specific mechanisms and parameters of the model, such as the cell population sizes in normal and treated conditions. However, only a few datasets are currently available for psoriasis. Additional bias could be attributed to decisions taken during the modeling process.

For instance, the model represents the proliferation only of keratinocytes. While the increase of immune cell populations due to their proliferation and recruitment is a characteristic of psoriasis, immune cell proliferation is not explicitly represented in the model, as the activation of T-helper cells is unrelated to their effector functions and cytokine secretion [51] we did not consider this necessary for the computational analysis of psoriasis development. Therefore, the population levels of keratinocytes and their physiological state (proliferation versus differentiation) were used as a proxy for the progress of the disease, considering that keratinocytes are the primary responders to disease progression or alleviation. Lastly, certain mechanisms that require a spatial description, such as cytokine diffusion or NETosis, cannot be explored in detail with the UPMaBoSS framework. To account for this, the model could be expanded to include such spatial properties and to better represent the spatial organization of the skin layers. The expanded model could be then analyzed as an agent based model with the PhysiBoSS framework [52]. With the current framework, we have demonstrated the potential of multiscale models of diseases to investigate the underlying disease pathogenesis and to identify potential targets for improving the efficacy of targeted therapies. In the future, the PsoriaSys model could be expanded to include other pertinent cell types and signaling pathways. An interesting and promising aspect, which could be explored through a dynamic Boolean model, involves the understanding of the development of tissue resident memory cells, a phenomenon associated with disease recurrence [53]. Moreover, enhancing the model with biomarkers and patient-specific data holds promise for predicting alternative paths of disease advancement, thereby offering insights into variation of treatment responses.

## Methods

### Model construction

A Prior Knowledge Network (PKN) encompassing the main inter- and intra-cellular signaling events taking place in the development of psoriasis was manually curated from scientific publications about the disease and its progression. The PKN consists of biological entities (nodes) with a reported role in the disease and their regulatory interactions (edges). Each node represents a cell type in various physiological states (called Cell nodes), a ligand or its respective receptors, a transcription factor, or other intermediate signaling proteins (collectively called Signaling nodes). The interactions between nodes were collected from the available literature and from databases with causal molecular interactions, such as SIGNOR 3.0 (Perfetto et al., 2016). Over decades of research on the disease, multiple innate and adaptive immune cells have been described for their role in psoriasis. However, the model focuses on the disease’s main cellular players, whose role is already well established.

Once the PKN was constructed, it was converted to a Boolean model. In Boolean models, the activity or state of a node can only take two possible values, 0 or 1. Nodes with a state of 0 are considered inactive, while active nodes take a value of 1. The state of a node is defined by its logical rules that define how the state of a node changes depending on the activity and the combinations of the nodes that regulate it. The logical rules were defined manually using the logical formalism AND, OR, and NOT. The general scheme for the definition of the logical rules for different node types is presented in Figure 4.

**Figure 4.**
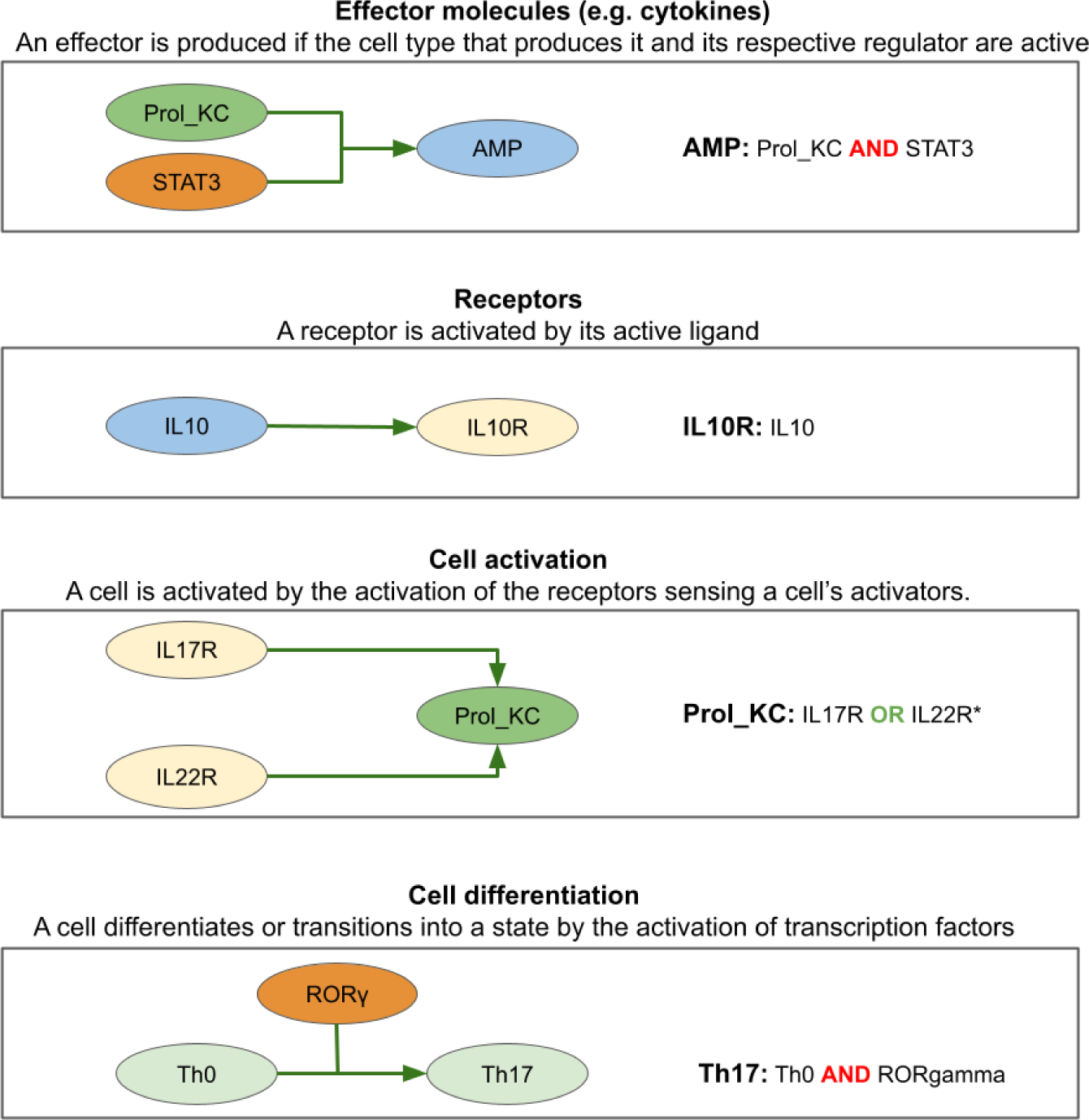
Schematic representation of the general scheme applied in the definition of logical rules in the PsoriaSys model.

### UPMaBoSS model

MaBoSS is based on Markov Chain processes applied on the Boolean network. A MaBoSS simulation computes trajectories of probabilities for nodes to be active over time. UPMaBoSS extends on the MaBoSS framework by adding dynamical properties such as the evolution of cell population size over time: cells can die, divide and communicate. The population update is done synchronously throughout the simulation (at every *t* step) until the maximum time is reached. The stochastic Boolean simulation framework UPMaBoSS [13] was used to approximate continuous behaviors, calculate the probabilities of node or phenotype activation, and estimate the cell population sizes.

First, to allow the simulation of population dynamics, two nodes representing ‘Division’ and ‘Death’ were added to the PsoriaSys model to allow the population dynamics simulation. The probabilities of activation are doubled for trajectories with Division node equal to 1, and for trajectories with the Death node equal to 1, the probabilities are set to 0. Each node of the model was assigned rates of activation and inactivation. When available, the parameters were defined based on available literature reporting on specifically psoriatic conditions or any other skin inflammatory states. As this type of measurement is scarce, only 17 parameters were defined based on prior knowledge (Supplementary Table 1). The remaining parameters were defined to be the same for nodes of the same type. For example, cytokine and chemokine ligands were set to undergo rapid degradation/inactivation (i.e., high inactivation rate), resulting from extracellular signal depletion and other mechanisms of inflammation control [16]. For receptors, the activation rates were updated in each step of the simulations based on the probability of their ligand being active; the more of a ligand produced, the higher the activation rate of its receptor. Due to the high number of parameters, a sensitivity analysis was performed to assess the robustness of the model and ensure that the assigned parameters were biologically reasonable. Each parameter was either reduced or increased by 50% of their initial value, similar to the sensitivity analysis performed in [12].

### Minimum Feedback Vertex Set (FVS) identification

The presence of feedback loops (or circuits) in a network has been shown to play a role in its dynamics: its ability to reach either multiple stable states in the presence of positive feedback loops or cyclic attractors in the presence of negative feedback loops [54]. The control of nodes that are part of such feedback loops (collectively called FVS) can be used to swift the network towards a target stable state. As a network can include numerous nodes that are part of the network’s FVS, a minimal set of nodes that can drive the system toward a desired state can be identified (minimum FVS). As this set of nodes is not unique, the intersection of three network propagation measures was used to rank 2-node combinations based on their ability to drive the model toward a desired state and, by extension, identify intervention points (i.e., drug targets) that would lead to psoriasis resolution. These topological measures included PRINCE propagation, Modified PRINCE, and CheiRank random walk, and their intersection is the highest performing for identifying the most influential minimum FVS [37]. The analysis was performed with a predefined minimum FVS size of two, so that the identified perturbations would reflect feasible interventions to treat psoriasis. The analysis requires Python 3.8. Instructions on creating a Conda virtual environment are included in the respective notebook.

## Data and code availability

The model and all scripts required for its analysis are provided as Jupyter Notebooks in https://github.com/Eirinits/PsoriaSys_model. The model is also submitted at BioModels, and can be accessed with the identifier MODEL2308300001.

## Author contribution

ET designed the project, built and analyzed the model. VN and LC aided and supervised the model building and analysis. MK supervised the project. ET drafted the manuscript with input from all authors. All authors have contributed and approved the manuscript.

## Acknowledgements

The authors would like to thank Dr. Mari Løset for her insights for the clinical phenotypes and control of psoriasis. Figure 1A has been generated with BioRender.

## Supplementary files

**Supplementary Table 1.**
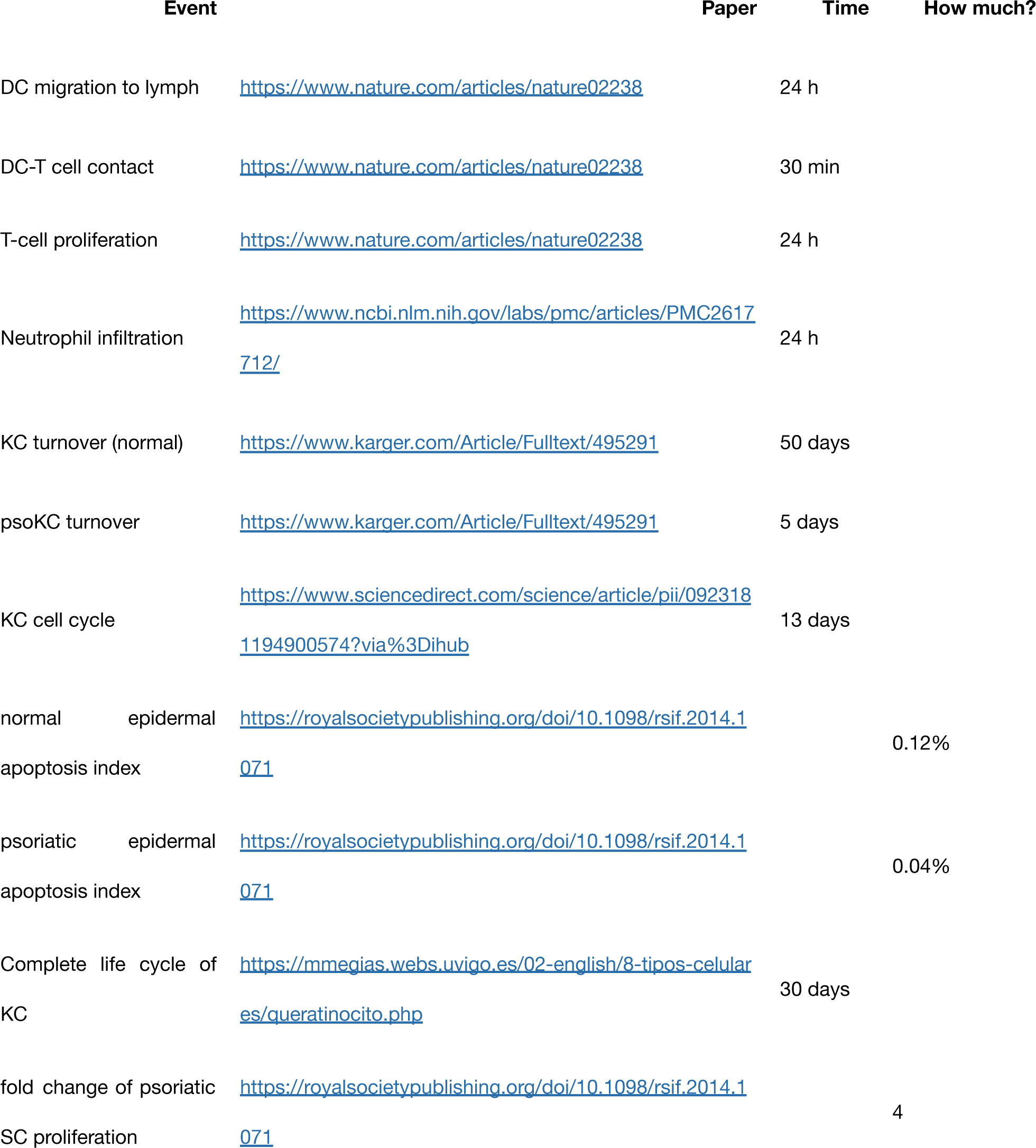
Rates and parameters.

**Supplementary Table 2.** The propagation consensus score is the score given by the consensus of three network propagation measures: PRINCE, modified PRINCE, and CheiRank.

| **Propagation**<br>**consensus score** | **Target 1** | **Target 2** |
| --- | --- | --- |
| 97 | STAT3 | Neutrophil |
| 95 | IL17 | Neutrophil |
| 95 | IL17R | STAT3 |
| 94 | STAT3 | PGE2 |
| 93 | IL17R | Neutrophil |
| 93 | IL17 | STAT3 |
| 92 | PGE2 | Neutrophil |
| 90 | TNFAR | STAT3 |
| 89 | STAT3 | TNFα |
| 88 | IL4 | Neutrophil |
| 87 | TNFα | Neutrophil |
| 86 | IFNγ | Neutrophil |
| 86 | STAT3 | IL4 |
| 86 | IL17 | PGE2 |
| 85 | IL17R | PGE2 |
| 84 | STAT3 | IL10 |
| 84 | STAT3 | IFNg |
| 84 | STAT3 | IL10R |
| 83 | TNFAR | Neutrophil |
| 82 | STAT3 | IFNGR |

**Supplementary Table 3.** Node reduction in all tested perturbations.

**Supplementary Figure 1.**
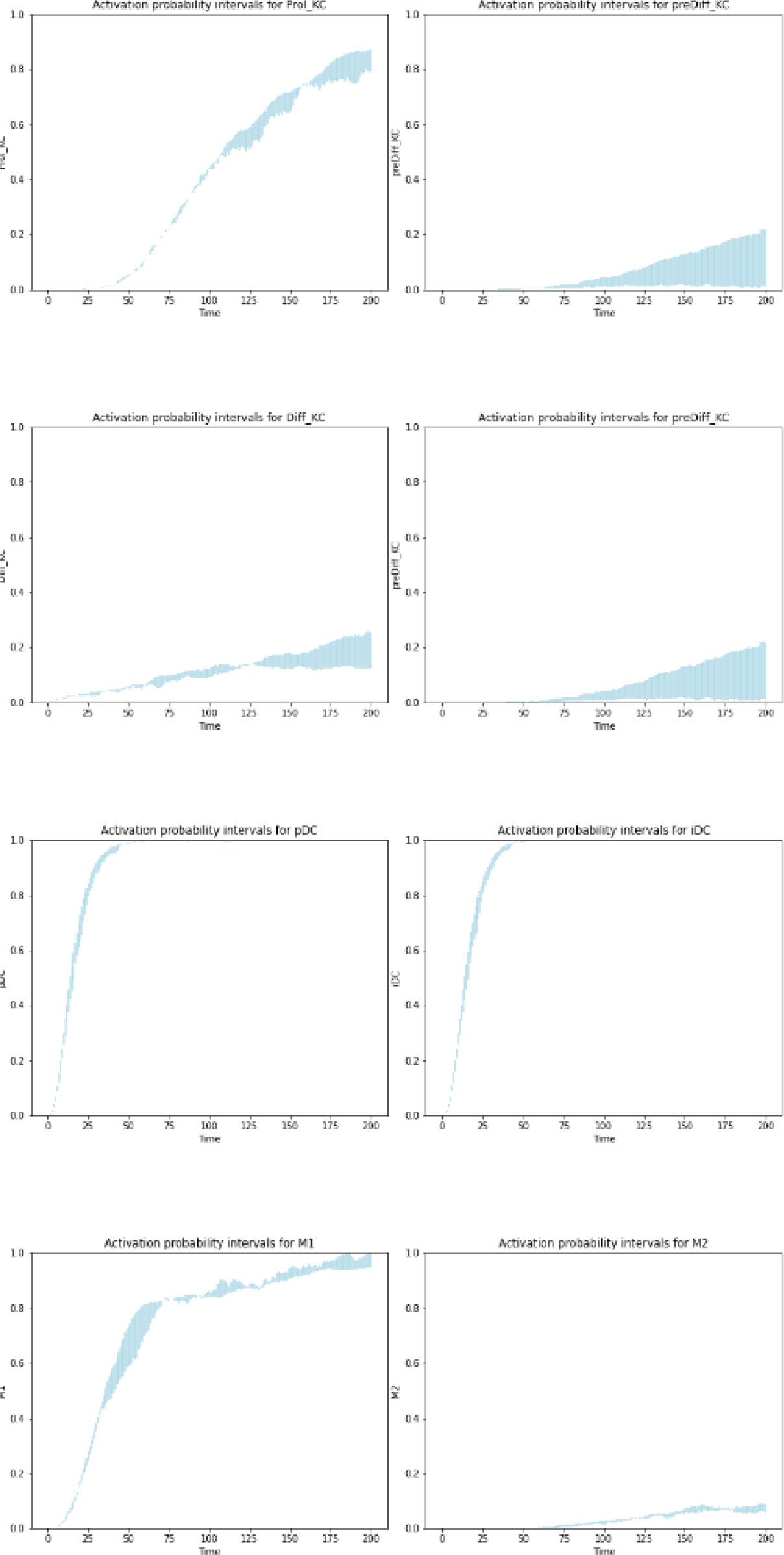
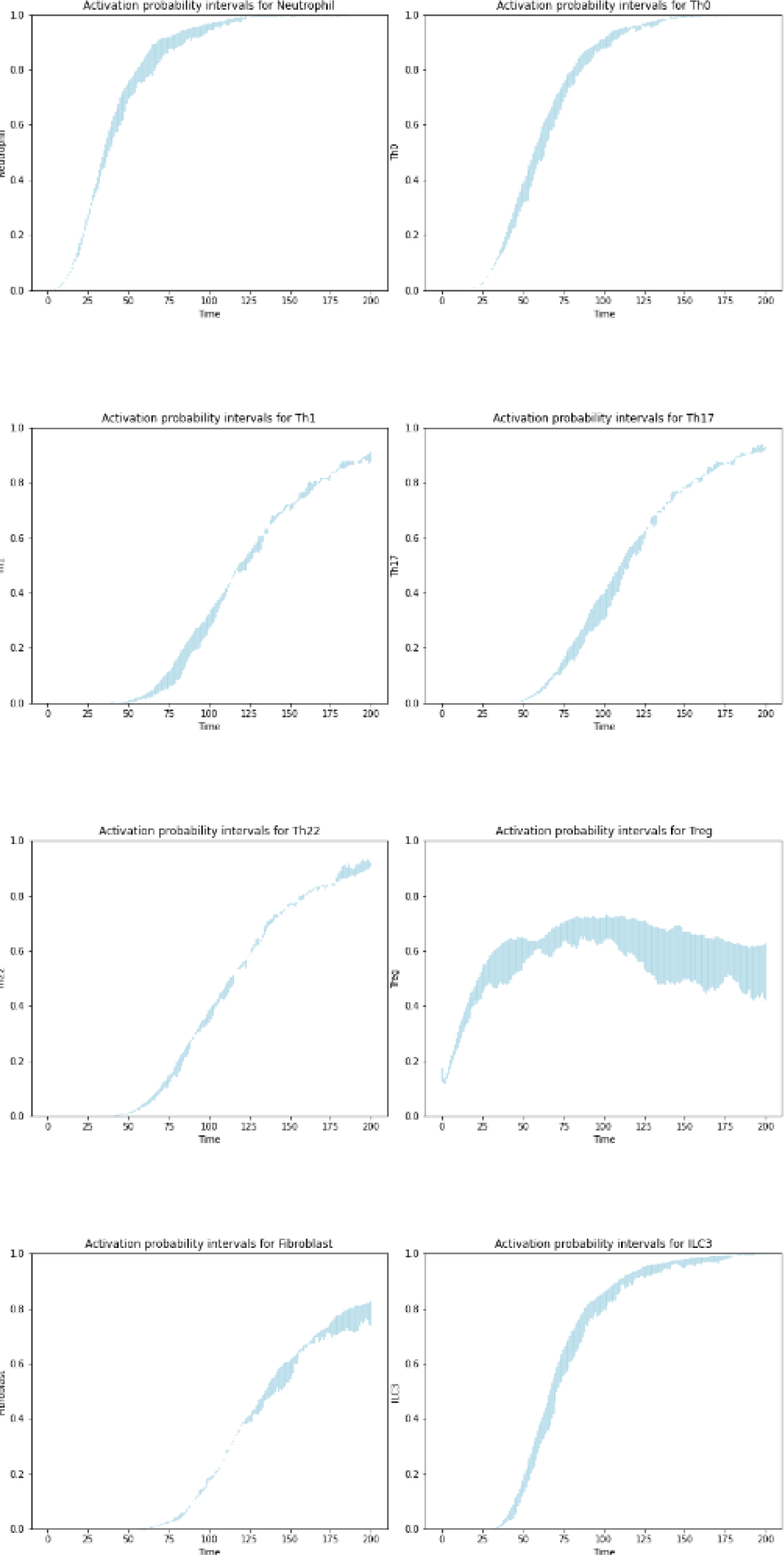
Sensitivity analysis results. Difference between the activation probability of each cell node between the wild-type model and model instances with adjusted activation/inactivation rates.

## Notes

### Competing Interest Statement

The authors have declared no competing interest.

## References

1. Kamiya K, Kishimoto M, Sugai J, Komine M, Ohtsuki M. Risk Factors for the Development of Psoriasis. Int J Mol Sci. 2019;20: 4347. doi:10.3390/ijms20184347

2. Boehncke W-H, Schön MP. Psoriasis. The Lancet. 2015;386: 983–994. doi:10.1016/S0140-6736(14)61909-7

3. Benhadou F, Mintoff D, del Marmol V. Psoriasis: Keratinocytes or Immune Cells – Which Is the Trigger? Dermatology. 2019;235: 91–100. doi:10.1159/000495291

4. Albanesi C, Madonna S, Gisondi P, Girolomoni G. The Interplay Between Keratinocytes and Immune Cells in the Pathogenesis of Psoriasis. Front Immunol. 2018;9. doi:10.3389/fimmu.2018.01549

5. Albanesi C. Immunology of Psoriasis. Clinical Immunology. Elsevier; 2019. pp. 871–878.e1. doi:10.1016/B978-0-7020-6896-6.00064-8

6. Wang A, Bai Y. Dendritic cells: The driver of psoriasis. J Dermatol. 2020;47: 104–113. doi:10.1111/1346-8138.15184

7. Casciano F, Pigatto PD, Secchiero P, Gambari R, Reali E. T Cell Hierarchy in the Pathogenesis of Psoriasis and Associated Cardiovascular Comorbidities. Front Immunol. 2018;9. doi:10.3389/fimmu.2018.01390

8. Al-Janabi A, Yiu ZZN. Biologics in Psoriasis: Updated Perspectives on Long-Term Safety and Risk Management. Psoriasis Targets Ther. 2022;12: 1–14. doi:10.2147/PTT.S328575

9. Damiani G, Odorici G, Pacifico A, Morrone A, Conic RRZ, Davidson T, et al. Secukinumab Loss of Efficacy Is Perfectly Counteracted by the Introduction of Combination Therapy (Rescue Therapy): Data from a Multicenter Real-Life Study in a Cohort of Italian Psoriatic Patients That Avoided Secukinumab Switching. Pharmaceuticals. 2022;15: 95. doi:10.3390/ph15010095

10. Diotallevi F, Paolinelli M, Radi G, Offidani A. Latest combination therapies in psoriasis: Narrative review of the literature. Dermatol Ther. 2022;35: e15759. doi:10.1111/dth.15759

11. Tsirvouli E, Aker E, Kuiper M. Patient-specific logical models replicate phenotype responses to psoriatic and anti-psoriatic stimuli. Systems Biology; 2023 Aug. doi:10.1101/2023.08.24.554583

12. Checcoli A, Pol JG, Naldi A, Noel V, Barillot E, Kroemer G, et al. Dynamical Boolean Modeling of Immunogenic Cell Death. Front Physiol. 2020;11. doi:10.3389/fphys.2020.590479

13. Stoll G, Naldi A, Noël V, Viara E, Barillot E, Kroemer G, et al. UPMaBoSS: A Novel Framework for Dynamic Cell Population Modeling. Front Mol Biosci. 2022;9. Available: https://www.frontiersin.org/articles/10.3389/fmolb.2022.800152

14. Stoll G, Caron B, Viara E, Dugourd A, Zinovyev A, Naldi A, et al. MaBoSS 2.0: an environment for stochastic Boolean modeling. Bioinformatics. 2017;33: 2226–2228. doi:10.1093/bioinformatics/btx123

15. Oza HB, Pandey R, Roper D, Al-Nuaimi Y, Spurgeon SK, Goodfellow M. Modelling and finite-time stability analysis of psoriasis pathogenesis. Int J Control. 2017;90: 1664–1677. doi:10.1080/00207179.2016.1217566

16. Sugimoto MA, Sousa LP, Pinho V, Perretti M, Teixeira MM. Resolution of Inflammation: What Controls Its Onset? Front Immunol. 2016;7. doi:10.3389/fimmu.2016.00160

17. Helm EY, Zhou L. Transcriptional regulation of innate lymphoid cells and T cells by aryl hydrocarbon receptor. Front Immunol. 2023;14. Available: https://www.frontiersin.org/articles/10.3389/fimmu.2023.1056267

18. Zheng SG. Regulatory T cells vs Th17: differentiation of Th17 versus Treg, are the mutually exclusive? Am J Clin Exp Immunol. 2013;2: 94–106.

19. Furue M, Furue K, Tsuji G, Nakahara T. Interleukin-17A and Keratinocytes in Psoriasis. Int J Mol Sci. 2020;21. doi:10.3390/ijms21041275

20. Brembilla NC, Senra L, Boehncke W-H. The IL-17 Family of Cytokines in Psoriasis: IL-17A and Beyond. Front Immunol. 2018;9. Available: https://www.frontiersin.org/articles/10.3389/fimmu.2018.01682

21. Mosca M, Hong J, Hadeler E, Hakimi M, Liao W, Bhutani T. The Role of IL-17 Cytokines in Psoriasis. ImmunoTargets Ther. 2021;10: 409–418. doi:10.2147/ITT.S240891

22. Blauvelt A, Chiricozzi A. The Immunologic Role of IL-17 in Psoriasis and Psoriatic Arthritis Pathogenesis. Clin Rev Allergy Immunol. 2018;55: 379–390. doi:10.1007/s12016-018-8702-3

23. Mylonas A, Conrad C. Psoriasis: Classical vs. Paradoxical. The Yin-Yang of TNF and Type I Interferon. Front Immunol. 2018;9. Available: https://www.frontiersin.org/articles/10.3389/fimmu.2018.02746

24. Chiricozzi A, Guttman-Yassky E, Suárez-Fariñas M, Nograles KE, Tian S, Cardinale I, et al. Integrative Responses to IL-17 and TNF-α in Human Keratinocytes Account for Key Inflammatory Pathogenic Circuits in Psoriasis. J Invest Dermatol. 2011;131: 677–687. doi:10.1038/jid.2010.340

25. Ten Bergen LL, Petrovic A, Krogh Aarebrot A, Appel S. The TNF/IL-23/IL-17 axis-Head-to-head trials comparing different biologics in psoriasis treatment. Scand J Immunol. 2020;92: e12946. doi:10.1111/sji.12946

26. Fania L, Morelli M, Scarponi C, Mercurio L, Scopelliti F, Cattani C, et al. Paradoxical psoriasis induced by TNF-α blockade shows immunological features typical of the early phase of psoriasis development. J Pathol Clin Res. 2020;6: 55–68. doi:10.1002/cjp2.147

27. Jadali Z. Unfulfilled Inflammatory Resolution: A Key Factor in the Pathogenesis of Psoriasis. Iran J Allergy Asthma Immunol. 2020;19: 337–347. doi:10.18502/ijaai.v19i4.4130

28. Roberts CA, Durham LE, Fleskens V, Evans HG, Taams LS. TNF Blockade Maintains an IL-10+ Phenotype in Human Effector CD4+ and CD8+ T Cells. Front Immunol. 2017;8. Available: https://www.frontiersin.org/articles/10.3389/fimmu.2017.00157

29. Wang B, Han D, Li F, Hou W, Wang L, Meng L, et al. Elevated IL-22 in psoriasis plays an anti-apoptotic role in keratinocytes through mediating Bcl-xL/Bax. Apoptosis. 2020;25: 663–673. doi:10.1007/s10495-020-01623-3

30. Jeon C, Sekhon S, Yan D, Afifi L, Nakamura M, Bhutani T. Monoclonal antibodies inhibiting IL-12, -23, and -17 for the treatment of psoriasis. Hum Vaccines Immunother. 2017;13: 2247–2259. doi:10.1080/21645515.2017.1356498

31. ten Bergen LL, Petrovic A, Krogh Aarebrot A, Appel S. The TNF/IL-23/IL-17 axis—Head-to-head trials comparing different biologics in psoriasis treatment. Scand J Immunol. 2020;92: e12946. doi:10.1111/sji.12946

32. Kim J, Bissonnette R, Lee J, Correa da Rosa J, Suárez-Fariñas M, Lowes MA, et al. The Spectrum of Mild to Severe Psoriasis Vulgaris Is Defined by a Common Activation of IL-17 Pathway Genes, but with Key Differences in Immune Regulatory Genes. J Invest Dermatol. 2016;136: 2173–2182. doi:10.1016/j.jid.2016.04.032

33. Tsirvouli E, Ashcroft F, Johansen B, Kuiper M. Logical and experimental modeling of cytokine and eicosanoid signaling in psoriatic keratinocytes. iScience. 2021;24: 103451. doi:10.1016/j.isci.2021.103451

34. Lee J, Aoki T, Thumkeo D, Siriwach R, Yao C, Narumiya S. T cell-intrinsic prostaglandin E2-EP2/EP4 signaling is critical in pathogenic TH17 cell-driven inflammation. J Allergy Clin Immunol. 2019;143: 631–643. doi:10.1016/j.jaci.2018.05.036

35. Al-Robaee AA, Al-Zolibani AA, Al-Shobili HA, Kazamel A, Settin A. IL-10 Implications in Psoriasis. Int J Health Sci. 2008;2: 53–58.

36. Ben Abdallah H, Johansen C, Iversen L. Key Signaling Pathways in Psoriasis: Recent Insights from Antipsoriatic Therapeutics. Psoriasis Targets Ther. 2021;11: 83–97. doi:10.2147/PTT.S294173

37. Newby E, Tejeda Zañudo JG, Albert R. Structure-based approach to identifying small sets of driver nodes in biological networks. Chaos Woodbury N. 2022;32: 063102. doi:10.1063/5.0080843

38. Chiang C-C, Cheng W-J, Korinek M, Lin C-Y, Hwang T-L. Neutrophils in Psoriasis. Front Immunol. 2019;10: 2376. doi:10.3389/fimmu.2019.02376

39. Zhang W, Dang E, Shi X, Jin L, Feng Z, Hu L, et al. The Pro-Inflammatory Cytokine IL-22 Up-Regulates Keratin 17 Expression in Keratinocytes via STAT3 and ERK1/2. PLOS ONE. 2012;7: e40797. doi:10.1371/journal.pone.0040797

40. Cai Y, Xue F, Quan C, Qu M, Liu N, Zhang Y, et al. A Critical Role of the IL-1β-IL-1R Signaling Pathway in Skin Inflammation and Psoriasis Pathogenesis. J Invest Dermatol. 2019;139: 146–156. doi:10.1016/j.jid.2018.07.025

41. Ghoreschi K, Thomas P, Breit S, Dugas M, Mailhammer R, van Eden W, et al. Interleukin-4 therapy of psoriasis induces Th2 responses and improves human autoimmune disease. Nat Med. 2003;9: 40–46. doi:10.1038/nm804

42. Hahn M, Ghoreschi K. The role of IL-4 in psoriasis. Expert Rev Clin Immunol. 2017;13: 171–173. doi:10.1080/1744666X.2017.1279054

43. Yousefzadeh H, Jabbari Azad F, Rastin M, Banihashemi M, Mahmoudi M. Expression of Th1 and Th2 Cytokine and Associated Transcription Factors in Peripheral Blood Mononuclear Cells and Correlation with Disease Severity. Rep Biochem Mol Biol. 2017;6: 102–111.

44. Liu P, He Y, Wang H, Kuang Y, Chen W, Li J, et al. The expression of mCTLA-4 in skin lesion inversely correlates with the severity of psoriasis. J Dermatol Sci. 2018;89: 233–240. doi:10.1016/j.jdermsci.2017.11.007

45. Chizzolini C, Brembilla NC. Prostaglandin E2: igniting the fire. Immunol Cell Biol. 2009;87: 510–511. 10.1038/icb.2009.56

46. Boniface K, Bak-Jensen KS, Li Y, Blumenschein WM, McGeachy MJ, McClanahan TK, et al. Prostaglandin E2 regulates Th17 cell differentiation and function through cyclic AMP and EP2/EP4 receptor signaling. J Exp Med. 2009;206: 535–548. doi:10.1084/jem.20082293

47. Ashcroft FJ, Mahammad N, Midtun Flatekvål H, J. Feuerherm A, Johansen B. cPLA2α Enzyme Inhibition Attenuates Inflammation and Keratinocyte Proliferation. Biomolecules. 2020;10: 1402. doi:10.3390/biom10101402

48. Yamamoto N, Koyama T, Nishino M, Iwasa S, Kondo S, Sudo K, et al. 534MO First in human study of ONO-4578, a PGE2-receptor EP4 antagonist, in monotherapy and combination with PD-1 checkpoint inhibitor nivolumab in patients with advanced or metastatic solid tumours. Ann Oncol. 2020;31: S468. doi:10.1016/j.annonc.2020.08.648

49. Rodriguez-Rosales YA, Langereis JD, Gorris MAJ, van den Reek JMPA, Fasse E, Netea MG, et al. Immunomodulatory aged neutrophils are augmented in blood and skin of psoriasis patients. J Allergy Clin Immunol. 2021;148: 1030–1040. doi:10.1016/j.jaci.2021.02.041

50. Zhou X, Chen Y, Cui L, Shi Y, Guo C. Advances in the pathogenesis of psoriasis: from keratinocyte perspective. Cell Death Dis. 2022;13: 1–13. doi:10.1038/s41419-022-04523-3

51. Brenchley JM, Douek DC, Ambrozak DR, Chatterji M, Betts MR, Davis LS, et al. Expansion of activated human naïve T-cells precedes effector function. Clin Exp Immunol. 2002;130: 431–440. doi:10.1046/j.1365-2249.2002.02015.x

52. Letort G, Montagud A, Stoll G, Heiland R, Barillot E, Macklin P, et al. PhysiBoSS: a multi-scale agent-based modelling framework integrating physical dimension and cell signalling. Bioinformatics. 2019;35: 1188–1196. doi:10.1093/bioinformatics/bty766

53. Owczarczyk-Saczonek A, Krajewska-Włodarczyk M, Kasprowicz-Furmańczyk M, Placek W. Immunological Memory of Psoriatic Lesions. Int J Mol Sci. 2020;21. doi:10.3390/ijms21020625

54. Thieffry D. Dynamical roles of biological regulatory circuits1. Brief Bioinform. 2007;8: 220–225. doi:10.1093/bib/bbm028

